# Mechanisms of Activation and Serotonin release from Human Enterochromaffin Cells

**DOI:** 10.1101/2025.03.17.643631

**Authors:** Constanza Alcaino, Nunzio Guccio, Emily L. Miedzybrodzka, Jaden R. Quale, Adam Davison, Christopher A. Smith, Emily Overington, Marta Santos-Hernández, Mae Tabbada, Tianyi Lu, Megan Hodge, Rula Bany-Bakar, Richard Kay, Ahmed Shaaban, Cordelia Imig, Frank Reimann, Fiona M. Gribble

## Abstract

**Background and Aims:** Gastrointestinal (GI) enterochromaffin (EC) cells are specialised sensors of luminal stimuli. They secrete most of the body’s serotonin (5-HT), and are critical for modulating GI motility, secretion, and sensation, while also signalling satiety and intestinal discomfort. The aim of this study was to investigate mechanisms underlying the regulation of human EC cells, and the relative importance of direct nutrient stimulation compared with neuronal and paracrine regulation.

**Methods:** Intestinal organoids from human duodenal biopsies were modified using CRISPR-Cas9 to specifically label EC cells with either the fluorescent protein Venus or the cAMP sensor Epac1-S. EC cells were purified by fluorescence-activated cell sorting for analysis by bulk RNA sequencing and liquid chromatography mass spectrometry peptidomics. The function of human EC cells was studied using single cell patch clamp, calcium and cAMP imaging and 5-HT ELISA assays.

**Results:** Human EC cells showed expression of receptors for nutrients (including *GPR142*, *GPBAR1, GPR119, FFAR2, OR51E1, OR51E2*), gut hormones (including *SSTR1,2&5*, *NPY1R, GIPR*) and neurotransmitters (*ADRA2A*, *ADRB1*). Functional assays revealed EC responses (calcium, cAMP and/or secretion) to a range of stimuli, including bacterial metabolites, aromatic amino acids and adrenergic agonists. Electrophysiological recordings showed that isovalerate increased action potential firing.

**Conclusions:** 5-HT release from EC cells controls many physiological functions and is currently being targeted to treat disorders of the gut-brain axis. Studying ECs from human organoids enables improved understanding of the molecular mechanisms underlying EC cell activation, which is fundamental for the development of new strategies to target 5-HT-related gut and metabolic disorders.

**Synopsis:** Human duodenal organoids expressing fluorescent proteins in enterochromaffin cells were used to study mechanisms underlying serotonin secretion. Different expression of key sensory receptors was identified by transcriptomic analysis, and validated by live cell second messenger imaging and secretion assays.

## INTRODUCTION

Enterochromaffin (EC) cells are the largest population of enteroendocrine cells (EECs) in the intestine and produce most of the body’s systemic serotonin (5-hydroxytryptamine or 5-HT). This 5-HT is critical for normal gastrointestinal function, modulating intestinal secretion, motility, and sensation, whilst also conveying a signal that modulates food intake^1^. Platelets take up 5-HT released from the gut and act as a high-capacity reservoir of 5-HT in the circulation, making it difficult to monitor acute release of 5-HT from EC cells in vivo, against this high background. Our understanding of the physiological stimuli controlling EC secretion, particularly in humans, is therefore relatively limited and mostly derived from studies on isolated cells and cell lines.

A number of studies have shown that EC cells are modulated by chemical and mechanical cues^2–4^, but it is debated whether small intestinal 5-HT secreting EC cells are directly nutrient-sensitive. A study in dogs reported elevation of 5-HT in portal blood after instillation of hypertonic glucose into the small intestine, but did not conclude whether it was glucose itself or the hypertonicity and consequent gut distension that acted as the trigger of 5-HT release^5^. Another study reported elevated circulating 5-HT levels in humans after ingestion of carbohydrates and fats, but suggested that the slow time course of the rise after the carbohydrate meal was more likely to reflect changes in peripheral 5-HT production or clearance than release from the gut^6^. Free sugars and fatty acids were, however, shown to trigger 5-HT secretion from intestinal biopsies and purified EC cell populations^7–10^. mRNA expression levels of nutrient-sensitive receptors in EC cells have been reported to be location-dependent and to vary between species^3, 8, 11, 12^ although one study suggested that small intestinal EC cells lack nutrient-sensing machinery and rely on paracrine signalling from L-cell derived Glucagon-like peptide 1(GLP-1)^11^.

Murine EC cells have also been shown to be responsive to irritant-activators of Trpa1 channels, to short and branched chain fatty acids (SCFA, BCFA) and to adrenergic receptor stimulation^3^. EC cell activation sensitizes sensory afferent neurons to mechanical stimulation and is critical for mechanically induced visceral hypersensitivity in mice^13^. In humans, diets low in FODMAPs (fermentable oligosaccharides, disaccharides, monosaccharides and polyols) reduce pain and diarrhoea in some patients with irritable bowel syndrome, potentially by reducing the action of microbially-produced SCFA on EC cells^14^. Germ-free mice have blunted expression of *Tph1* (tryptophan hydroxylase – a key step in 5-HT biosynthesis in EC cells) and colonic 5-HT content^15^, suggesting that microbiota-derived metabolites such as SCFAs can also modulate EC cell numbers or expression profiles^16,17^.

The extent to which EC cell secretion is regulated by local paracrine signals remains unclear, although candidate local paracrine modulators have been suggested previously. Somatostatin (SST), which inhibits other EEC subtypes such as L-cells^18^, also reduced 5-HT release from the carcinoid cell line BON, often used as an EC model cell line^19^. In the colon, a few studies have shown that insulin-like peptide 5 (INSL5), a hormone co-secreted from L-cells, stimulates colonic motility through a pathway sensitive to 5-HT_3_ receptor inhibition, suggesting potential cross-talk with local EC cells; however, whilst the INSL5 receptor Rxfp4 is expressed by mouse EC cells, the preferential G_i_-coupling of this receptor would suggest inhibition rather than stimulation of 5-HT secretion from EC-cells.^20, 21^.

In this study, we aimed to characterize the mechanisms underlying 5-HT secretion from human EC cells by developing transgenic EC-labelled organoids derived from duodenal biopsies, using CRISPR-Cas9 homology-directed repair (HDR) to express fluorescent proteins specifically in cells expressing *TPH1*. Fluorescently labelled EC cells from these organoids were purified by flow cytometry for bulk RNA sequencing and peptidomic analysis and studied by live cell fluorescent imaging to examine functional responses to stimuli such as nutrients, neurotransmitters and hormones, and by electrophysiology to monitor action potential firing and vesicular release as changes in cell capacitance. This work provides the first functional characterisation of primary (non-tumour-derived) human EC cells, which is critical for our understanding of their role in disorders of the gut-brain axis.

## METHODS

### Human organoid culture and maintenance

Human duodenal organoids were derived from anonymous surgical samples from Addenbrooke’s Hospital Tissue Bank (Cambridge, UK), under ethical approval by East of England–Cambridge Central Research Ethics Committee (license number 09/H0308/24). Duodenal organoids were generated and maintained as described previously^22^. Cultures were maintained as domes of several 3D spheroids embedded in basement membrane extract (BME) on the base of tissue culture plates. Cultures were passaged every 7 days by adding 1-4 mature domes to TrypLE (Thermofisher) at 37°C for 7 min and triturating via pipetting through a P1000 pipette 20-30 times. Residual small cell clusters were then embedded in BME and seeded onto prewarmed plates. To promote differentiation, EGF was removed from the culture medium 2-3 days after seeding (IF medium). Subsequently, organoids were grown in IF medium for 5-7 days, at which point fluorescent TPH1-Venus cells could be observed. For access to the Venus-positive cells for imaging and electrophysiological experiments, 2-D monolayer cultures were made. For this, organoids were dissociated as described and seeded onto 2% Matrigel (Corning) coated 35mm plastic or glass coverslips and glass bottom dishes for electrophysiology and imaging experiments respectively and incubated overnight.

### Generation of TPH1-Venus human duodenal organoids

CRISPR-Cas9 homology directed repair (HDR) was used to knock in either Venus or the FRET cAMP reporter EpacS-H187 ^23^ following a P2A sequence to enable bicistronic expression under control of the TPH1 promoter (Figure 1A). A CRISPR site (ACTGGCTACTGTTAGATACTCGG) close to the stop codon (underlined) in exon 10 was targeted. SgRNA-Cas9 and donor plasmids were generated, purified, and prepared for electroporation, as described previously ^24, 25^. Successful recombinants were enriched by antibiotic selection by adding G418 (0.5 mg/ml) to the medium 3-7 days after electroporation. Surviving organoids were manually picked and seeded in single BME domes to ensure establishment of monoclonal organoid lines. DNA was extracted from each organoid using QuickExtract DNA Extraction Solution (Lucigen Corporation, USA) and successful integration was assessed by PCR genotyping and confirmed by Sanger sequencing (Source Bioscience).

**Figure 1.**
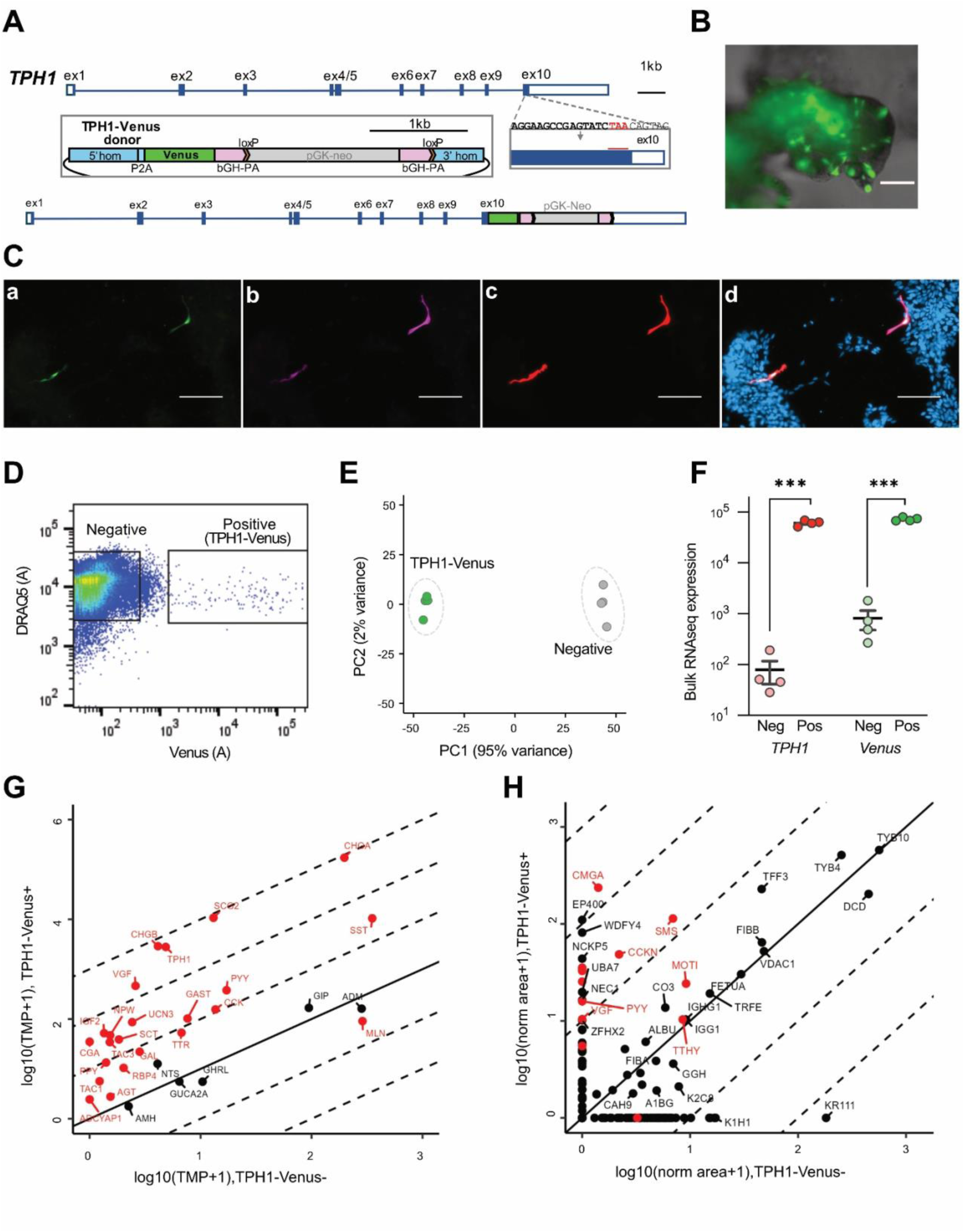
Development and expression profile of human duodenal Enterochromaffin (EC) cells. A) Schematic showing the knock-in strategy to insert yellow fluorescent protein venus gene in exon 10 of the *TPH1* gene using CRISPR-Cas9 homology donor repair. The same strategy was used to insert an Epac1-S-H187 sensor for cAMP measurements. B) Image of duodenal 3D organoid from TPH1-Venus line. Venus+ cells are shown in green. Scale bar represents 400 µm. C) Immunocytochemistry of fixed TPH1-Venus human duodenal 2D epithelial monolayer cultures. Cells staining positive for (a) GFP/Venus (*green*) are also immunopositive for (b) 5-HT (*magenta*) and (c) Chromogranin A (*red*). (d) Merged image of a-c with DAPI nuclear staining. Scale bars represent 100 µm. D) Representative fluorescent-activated cell sorting (FACS) plot. TPH1-Venus positive and negative cells were isolated based on Venus fluorescence intensity, after selection for live DAPI-negative and DRAQ5-positive cells only. E) Principal component analysis (PCA) plot showing TPH1-Venus positive (green) and negative (grey) cell populations following bulk RNA sequencing analysis. F) Expression of *TPH1* and *Venus* genes in sorted Venus-positive and -negative populations following bulk RNA sequencing analysis. G) Transcriptomic analysis plot showing expression of genes of interest in TPH1-Venus positive (y-axis) versus TPH1-Venus negative populations (x-axis). H) LC-MS/MS peptidomic analysis of purified TPH1-Venus positive and negative cells (the individual peptides detected are combined and associated to the parental protein, labelled by protein name and expressed as mean peak area). Proteins of interest are highlighted in red.

### cDNA library preparation and RNA sequencing

RNA-extraction and sequencing were performed after fluorescence-activated cell sorting (FACS) as previously described ^24, 25^. Gene expression is presented in transcripts per million. Differential expression was calculated using the Wald test (default in DESeq2), comparing TPH1-Venus positive vs negative cell populations.

### Peptidomic analysis

Peptide-extraction and analysis of FACS sorted cells were performed by LC-MS/MS as previously described ^24, 25^.

### Secretion assays

Differentiated organoids were removed from domes using ice-cold ADF and centrifuged at 400g for 4 min. Organoids were washed twice for 15 minutes at 37°C with saline buffer supplemented with 1 mM glucose, 0.1% bovine serum albumin (BSA), 10 μM fluoxetine and 10 μM 5-HT stabiliser (from kit). Approximately 30 organoids per well were distributed into V-bottom 96-well plates and incubated in duplicate or triplicate wells with test reagents (or vehicle control) dissolved in saline buffer for 1h at 37°C. Subsequently plates were centrifuged at 2000g for 5 min at 4°C, and supernatants collected into individual tubes were snap-frozen prior to analysis. Total 5-HT levels in the supernatants were measured by LDN Serotonin ELISA kit.

### Calcium and cAMP imaging

Live single-cell calcium imaging of Venus-expressing EC-cells was performed as previously described after loading with Fura2-AM ^24–26^. cAMP-dependent FRET imaging was performed on TPH1-Epac-S-H187 organoids as described previously^22^.

### Electrophysiology

Patch clamp recordings of individual Venus-fluorescent cells were performed using 2D monolayer cultures 1-2 days after seeding, at room temperature in extracellular saline solution. Patch pipettes were pulled from borosilicate glass (1.5 mm OD, 1.17 mm ID, Harvard Apparatus) to 3-7 MΩ using either a p-2000 puller (Sutter Instruments) or a PC-100 puller (Narishige, Japan) and fire polished using a microforge. Tips were coated with refined yellow beeswax for perforated patch clamp experiments.

Perforated patch-clamp recordings were performed using an Axopatch 200B amplifier connected through a Digidata 1440A A/D converter and visualised using pCLAMP software (Molecular Devices). Pipettes were filled with an intracellular solution consisting of (mmol/l): 76 K_2_S0_4_, 10 NaCl, 10 KCl, 10 HEPES, 55 sucrose, 1 MgCl_2_; adjusted to pH 7.2 with KOH, and the extracellular solution consisted of (mmol/l) 138 NaCl, 4.5 KCl, 10 HEPES, 4.2 NaHCO_3_, 2.6 CaCl_2_, 1.2 MgCl_2_, 1.2 Na_2_HPO_4_, 1 D-Glucose. A stock amphotericin-B solution dissolved in DMSO was prepared fresh on the day of recording and diluted in intracellular solution to a final concentration of 200 µg/ml. Evoked action potentials were recorded in current-clamp mode by applying square current steps of increasing size (50 or 500ms, Δ2-30 pA) from a baseline potential of −70mV.

Whole-cell voltage clamp capacitance recordings were performed using the Sine ± DC technique in the lockin extension of the HEKA EPC-10 amplifier and Patchmaster software (HEKA). For measurements of pool size, the intracellular solution consisted of (mmol/l): 123 Glutamic Acid, 40 Cs-HEPES, 2 Mg-ATP, 1 MgCl_2_, 17 NaCl, 0.26 EGTA, adjusted to pH 7.2 with CsOH and the extracellular solution consisted of (mmol/l) 145 NaCl, 2.8 KCl, 2 CaCl_2_, 1 MgCl_2_, 10 HEPES, 11 glucose. Osmolarity was adjusted to 300-310 mOsm. Pool size measurements were performed using a 10-pulse train protocol of 6 short depolarisations followed by four longer depolarisations (−70 to +20mV, 10 and 100ms respectively). For capacitance measurements of potentiation of exocytosis, the intracellular solution consisted of (mmol/l) 125 Caesium Gluconate, 10 TEA-Cl, 10 NaCl, 1 MgCl_2_, 5 HEPES, 10 EGTA, 3 Mg-ATP, 0.5 GTP, adjusted to pH 7.2 with CsOH and the extracellular solution consisted of (mmol/l) 138 NaCl, 4.5 KCl, 10 HEPES, 4.2 NaHCO_3_, 2.6 CaCl_2_, 1.2 MgCl_2_, 1.2 Na_2_HPO_4_, 1 D-Glucose, supplemented with 200 µmol/l Isovalerate or 10 μmol/l of Forskolin and 100 μmol/l of IBMX dependent on recording condition. Capacitance recordings were performed using a 15-pulse train of square depolarisations (−80 to +0mV, 500ms). Analysis of capacitance recordings was performed using either Igor Pro 8 (Wavemetrics) or RStudio (Posit), where capacitance measurements were taken as the average of the capacitance response in the 300ms interstimulus interval.

For the experiments assessing different vesicular pools, the first six 10 ms pulses should primarily activate the IRP, as only a very brief influx of Ca^2+^ is induced during these pulses. The last four 100 ms pulses allow sustained Ca^2+^ entry which triggers fusion of the RRP. Fusion of vesicles with the plasma membrane increases membrane surface area^27^, which can be recorded as changes in the total membrane capacitance, with the capacitance change reflecting net vesicular fusion (exocytosis minus endocytosis). Out of the 23 total human EC-cells measured only five made it into the final analysis; 9 cells (39%) were excluded because they exhibited no exocytosis (defined by a lack of capacitance change despite the presence of robust inward currents), and 3 (13%) were excluded because they exhibited spontaneous exocytosis (defined by capacitance measurements that fluctuated without correlation to current traces). Of the remaining 11 cells that exhibited provoked exocytosis (defined by a clear capacitance increase following current peaks), 2 were excluded due to unstable patch measurements, and 4 were excluded because they required more than one application of the depolarisation protocol before exocytosis was evident.

### Immunocytochemistry

Antibodies used were as follows; rabbit αGFP (1:1000, Nordic Biosite), goat α5-HT (1:1000, Immunostar), rabbit αCgA (1:50, Abcam), DαR555 (1:1000, ThermoFisher Scientific), DαG633 (1:1000, ThermoFisher Scientific). Immunocytochemistry was performed on 2D monolayer cultures of human TPH1-venus organoids which were prepared as described above. 48 hrs post-seeding cultures were washed 3×5 minutes in PBS and fixed in 4% paraformaldehyde for 30 minutes. All washes and incubations were performed at room temperature in a rocker set to 12 rpm. Cultures were then washed 3×5 minutes in PBS and left in blocking solution (10% serum, 0.3% Triton X-100) for 45 minutes then incubated in antibody solution containing primary antibodies (5% serum, 0.1% Triton X-100) for 2 hours. Cultures were then washed 3 times in PBS and incubated in antibody solution containing secondary antibodies for 1 hour. Cultures were washed a further 3 times in PBS and then incubated in DAPI solution (0.1% DAPI in PBS) for 10 minutes, washed a further 3 times in PBS and mounted using Aqua-Poly/Mount mounting media. Slides were imaged using a Zeiss Axio Observer microscope with an Axiocam 702 camera and a Colibri 5/7 LED light source within the ZEN Microscopy software (ZEISS), using consistent intensities and exposure times across samples.

### Buffers

Secretion assays, live fluorescent microscopy (Ca^2+^ and cAMP measurements) and electrophysiological experiments were carried out using saline buffer containing (in mmol/l): 138 NaCl, 4.5 KCl, 4.2 NaHCO_3_, 1.2 NaH_2_PO_4_, 2.6 CaCl_2_, 1.2 MgCl_2_, 10 HEPES; adjusted to pH 7.4 with NaOH. Glucose and test compounds were added as indicated.

### Data analysis

Calcium and cAMP data were not normally distributed, so expressed as median (IQR). Secretion data are presented as mean +/− SEM. Statistical tests were performed using GraphPad Prism v10, DESeq2 (RNA sequencing) or R v12. For calcium and cAMP imaging experiments, cells were classified as ‘responders’ if the *z* score was >3 for at least two consecutive timepoints during perfusion of test substance; *z* score = [(*F_t_* − mean *F*_b_)/SD *F*_b_], where *F_t_* is the ratio (fura2 or FRET) at timepoint *t* during perfusion of the test reagent, *F_b_* is the ratio under basal conditions, and mean *F_b_* and SD *F_b_* are calculated from 60 s of datapoints prior to test addition. All cells that had a positive z-score for their positive control (KCl or Forskolin+IBMX) were included in further statistical analyses. For statistical determination of whether there was a significant calcium or cAMP response across all cells tested with a specific agent, Fura2 or FRET ratios in the presence of the test agent were normalised to the ratio under basal conditions measured for each cell. Statistically significant elevations above baseline were determined using a one-sample Wilcoxon test (against a baseline of 1). Inhibitory effects of agents on cAMP in the presence of IBMX were assessed by Friedman test with post hoc Dunn’s multiple comparisons test. 5-HT secretion was normalised to the mean basal measurement in the same plate and significance tested by one-sample t-test against a baseline of 1.

## RESULTS

### Development and characterisation of TPH1-Venus+ EC cells from human duodenal organoids

To selectively label 5-HT-releasing human EC cells we generated duodenal organoids from patient-derived biopsies and used CRISPR-Cas9-mediated homology directed repair to express the yellow fluorescent protein Venus under the control of the *TPH1* promoter (Figure 1A). Resultant TPH1-Venus organoids exhibited sporadic green fluorescent cells, with a pattern and morphology consistent with fluorescent labelling of EC cells (Figure 1B). To validate the model and assess the morphology of human organoid-derived EC cells, 2D monolayer cultures from TPH1-Venus organoids were immunostained using antibodies against Green fluorescent protein/Venus, 5-HT and Chromogranin A (Figure 1C), which confirmed specific expression of Venus in cells staining positive for 5-HT and chromogranin A.

Venus positive and negative cells were purified by fluorescence-activated cell sorting from organoid cultures, for bulk RNA sequencing and mass spectrometry peptidomics (Figure 1D-H). Principal component analysis of the RNA-seq results showed clear separation of the fluorescent and non-fluorescent cell populations (Figure 1E). The expression of both *TPH1* and *Venus* were significantly higher in the *TPH1*-Venus+ fraction, compared with non-fluorescent cells, further validating that fluorescent reporter marked EC cells (Figure 1F). In addition to *TPH1*, the fluorescently labelled cells were enriched for the expression of other gut hormones, such as the expected *TAC1*, a known EC cell peptide precursor, and also for peptide YY (*PYY)*, *SST* and cholecystokinin (*CCK*) (Figure 1G). Functional translation of these gut hormone mRNAs was confirmed by peptidomic mass spectrometry analysis of purified EC cells, which identified enriched production of Chromogranin A, CCK, PYY and SST (Figure 1H).

### Expression of Neurotransmitter Receptors in Human Enterochromaffin Cells

Human TPH1-Venus (EC) cells expressed a variety of neurotransmitter receptors, including GABA, dopamine, acetylcholine, and adrenergic receptors (Supplementary Figure 1), suggesting modulation of EC cells by both intrinsic and extrinsic neurons found in the gut epithelium, consistent with previous reports^3, 28^. Contrasting with the previously reported expression profile of mouse *ChgA*-positive cells, however, which identified a lack of beta-adrenergic receptors^3^, human EC cells exhibited expression of *ADRA2A, ADRA2B, and ADRB1,* of which *ADRA2B* and *ADRB1* were enriched in fluorescent compared with non-fluorescent cells (Figure 2A). Dopamine receptors, known to also be partially activated by adrenergic agonists, were also expressed in human EC cells, with *DRD2* notably enriched in the TPH1-fluorescent population.

**Figure 2.**
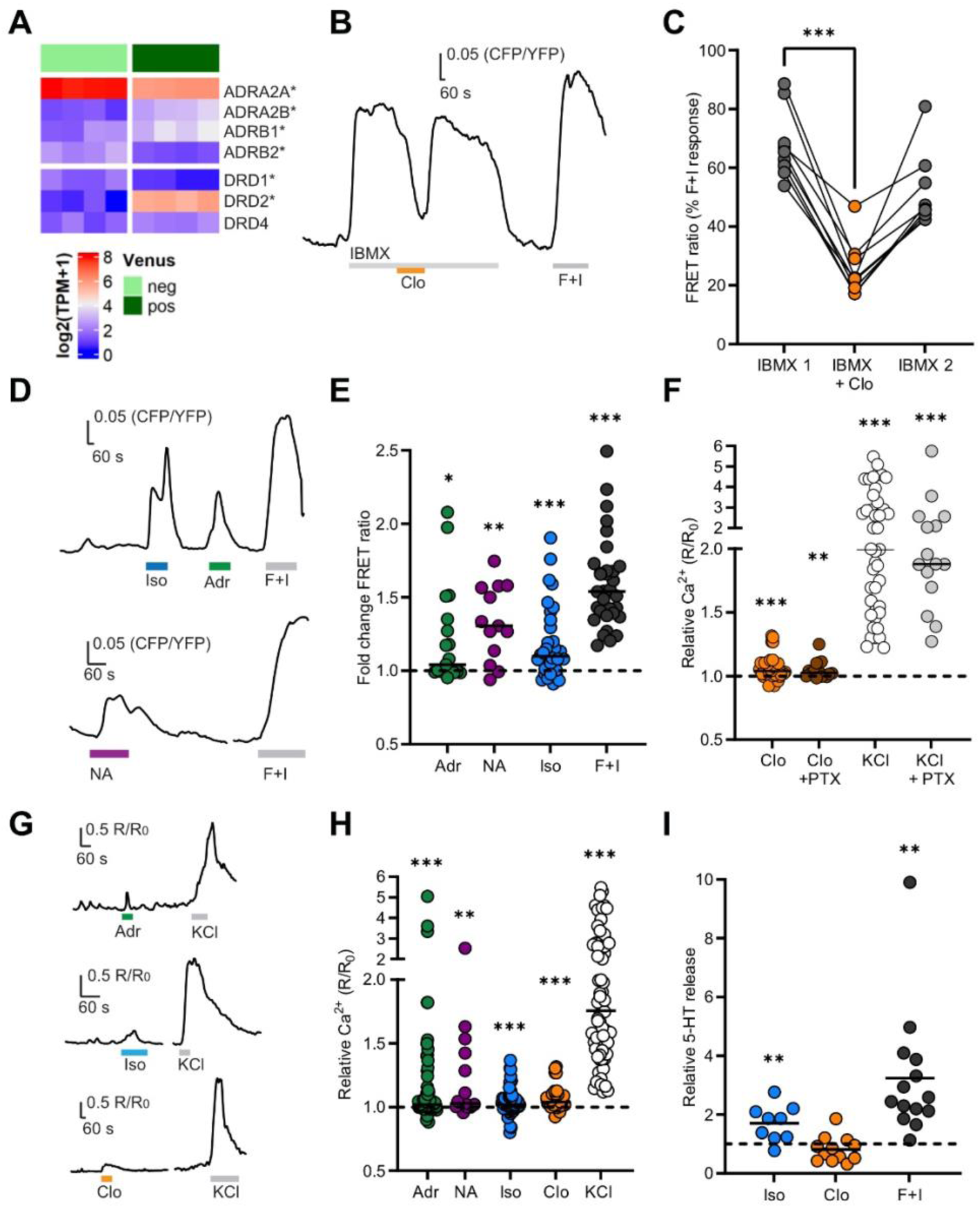
Adrenergic activation of human Enterochromaffin cells. A) Heatmap showing expression of adrenergic and dopaminergic receptors in TPH1-Venus-positive (dark green) and -negative (clear green) cell populations. * Denotes significant enrichment in Venus+ population by Wald test (p < 0.05). B) Representative FRET (CFP/YFP) ratio trace for single EC cell perfused with a pre-incubation of IBMX (100 μM), followed by Clonidine (Clo, 1 µM) and positive control Forskolin + IBMX (10 μM and 100 μM, respectively; F+I). C) Mean data collected as in (B), shown as the maximum CFP/YFP ratios during perfusion of IBMX1 (before clonidine addition) and IBMX2 (after clonidine washout) and the minimum CFP/YPF ratio during perfusion of IBMX + Clonidine (Clo), all expressed as a percentage of the response to F+I recorded in the same cell. *** p<0.001 by Friedman test with post hoc Dunn’s multiple comparisons test. D) Representative FRET (CFP/YFP) ratio traces of single EC cells being perfused with (E) Isoproterenol (Iso, 1 μM), Adrenaline (Adr, 30 μM), Noradrenaline (NA, 30 μM) and Forskolin + IBMX (F+I, 10 μM and 100 μM, respectively). E). Fold change of FRET ratio across different cells recorded as in (D). Data are shown as ratio between maximal CFP/YFP ratio (R) during perfusion of stimulus and maximal CFP/YFP ratio (R_0_) during perfusion of basal solution. (n = 13-34 cells from 2 to 4 independent experiments). * p<0.05, ** p<0.01, *** p<0.001 by one sample Wilcoxon test. F) Mean data showing the fold change in Fura2 ratio R/R_0_ during perfusion of clonidine (Clo, 1 µM) or KCl (70 mM), following pre-incubation for 24 hours at 37C with either control media or media treated with 200 ng/mL pertussis toxin (PTX). p=0.39 for comparison between clonidine and clonidine+PTX by Mann-Whitney test. G) Representative Fura2 (340/380 nm) ratio traces showing responses of single EC cells perfused with adrenaline (Adr, 30 μM), isoproterenol (Iso, 1 μM) and clonidine (Clo, 1 μM), with KCl (70 mM) as a positive control. H) Combined data from multiple cells recorded as in (G), or with noradrenaline (NA, 30 μM), shown as ratio between R (Fura2 ratio during perfusion of stimulus) and R_0_ (Fura2 ratio during perfusion of basal solution). (n = 16-72 cells from independent experiments). ** p<0.01, *** p<0.001 by one sample Wilcoxon test. I) Fold change of supernatant 5-HT concentrations in human duodenal TPH1-Venus positive 3D organoids incubated for 1 hour with isoproterenol (Iso,1 μM), clonidine (Clo,1 μM) and forskolin + IBMX (F+I, 10 μM and 100 μM, respectively). Results are presented relative to concentrations measured in control wells from the same plate. ** p<0.01 by one sample t-test.

To understand the role of cAMP following the activation of Gs- and Gi-coupled adrenergic receptors, we developed a TPH1-Epac1-S organoid line to measure changes in intracellular cAMP concentrations using fluorescence resonance energy transfer (FRET) imaging in 2D organoid cultures, as previously reported in a GIP-Epac1-S duodenal organoid line^22^, enabling use of the CFP/YFP FRET ratio as a measure of the intracellular cAMP concentration (Figure 2B-E). The selective ADRA2 agonist clonidine (1 µM) reduced cAMP levels in EC cells in the presence of the non-selective phosphodiesterase inhibitor 3-isobutyl-1-methylxanthine (IBMX, 100 μM) (Figure 2B,C); whereas application of the ADRB1 agonist isoproterenol (1 µM) increased cAMP levels (median 1.10-fold (IQR 1.00-1.31) FRET change; p<0.001) (Figure 2D,E). The non-selective adrenoreceptor agonists adrenaline (30 μM) and noradrenaline (30 μM) also significantly elevated EC cell cAMP levels (FRET changes of 1.04-fold (1.00-1.35), p=0.036; 1.31-fold (1.09-1.57), p=0.002; respectively) (Figure 2D,E).

A previous study on murine EC cells demonstrated that activation of the Gi-coupled Adra2a could increase intracellular Ca^2+^ concentrations by opening the transient receptor potential channel Trpc4. We tested the response to ADRA2 activation in human EC cells using the ratiometric Ca^2+^ dye Fura2-AM, revealing that even though clonidine (1 µM) decreased cAMP, it induced a small but significant increase in the Fura2 ratio, indicative of an increase in the intracellular Ca^2+^ concentration (median 1.04 (IQR 1.02-1.10) Fura2 R/R_0_, p<0.001; 23/42 cells exhibited significant responses by z-score analysis). Significant clonidine responses were also observed after blocking Gαi-coupling with pertussis toxin (PTX) (1.03 (1.00-1.10) Fura2 R/R_0_, p=0.004; 10/15 cells responded) (Figure 2F). EC cell Ca^2+^ responses were additionally observed in response to adrenaline (30 μM; 1.02 (1.00-1.16) Fura2 R/R_0_, p<0.001; 19/44 cells responded), noradrenaline (30 μM; 1.03 (1.01-1.36) Fura2 R/R_0_, p=0.001; 11/19 cells responded) and isoproterenol (1 µM; 1.02 (0.99-1.07) Fura2 R/R_0_, p<0.001; 14/38 cells responded) (Figure 2G,H).

To study the effects of adrenergic receptor agonists on 5-HT release from human EC cells, we performed secretion assays on 3D organoids and measured 5-HT release by ELISA after 30 min test incubations. Isoproterenol (1 µM), but not clonidine (1 µM), elicited a significant increase in 5-HT release compared with baseline (Figure 2I) (1.7±0.2 fold, p=0.009; and 0.8±0.1 fold, p=0.18; respectively). Overall, these data suggest that human EC cell activation by adrenergic agonists relies heavily on ADRB1, contrasting with the report of a predominant role for Adra2 in murine EC-cells^3, 13^.

### Paracrine Activation of Human Enterochromaffin Cells by Gut Hormones

We found differential expression of several gut hormone receptors in human EC cells, including the Gi-coupled SST receptors *SSTRs 1, 2, 3 and 5* and the PYY receptor *NPY1R*, the Gq-coupled CCK receptor *CCKAR* and the Gs-coupled incretin hormone receptor *GIPR* (Figure 3A). To our surprise, and despite INSL5 production being restricted to the distal colon and rectum, we found enriched expression of *RXFP4* in duodenal EC cells (Figure 3A). Activation of GIPR (using 100 nM GIP(1-42)) triggered significant cAMP responses in EC cells (FRET change 1.08 (1.04-1.20), p<0.001) (Figure 3B,C). GIP (100nM) also significantly stimulated 5-HT release (1.5±0.2 fold above baseline; p=0.035; Figure 3D). CCK (100 nM) induced a small, but significant Ca^2+^ response in 6/10 cells (1.02 (1.00-1.07) Fura2 R/R_0_, p=0.042) (Figure 3D,E), possibly reflecting the relatively low *CCKAR* expression level. Activation of Gi-coupled receptors by SST, PYY and INSL5 lowered EC cell cAMP levels (Figure 3G-L).

**Figure 3.**
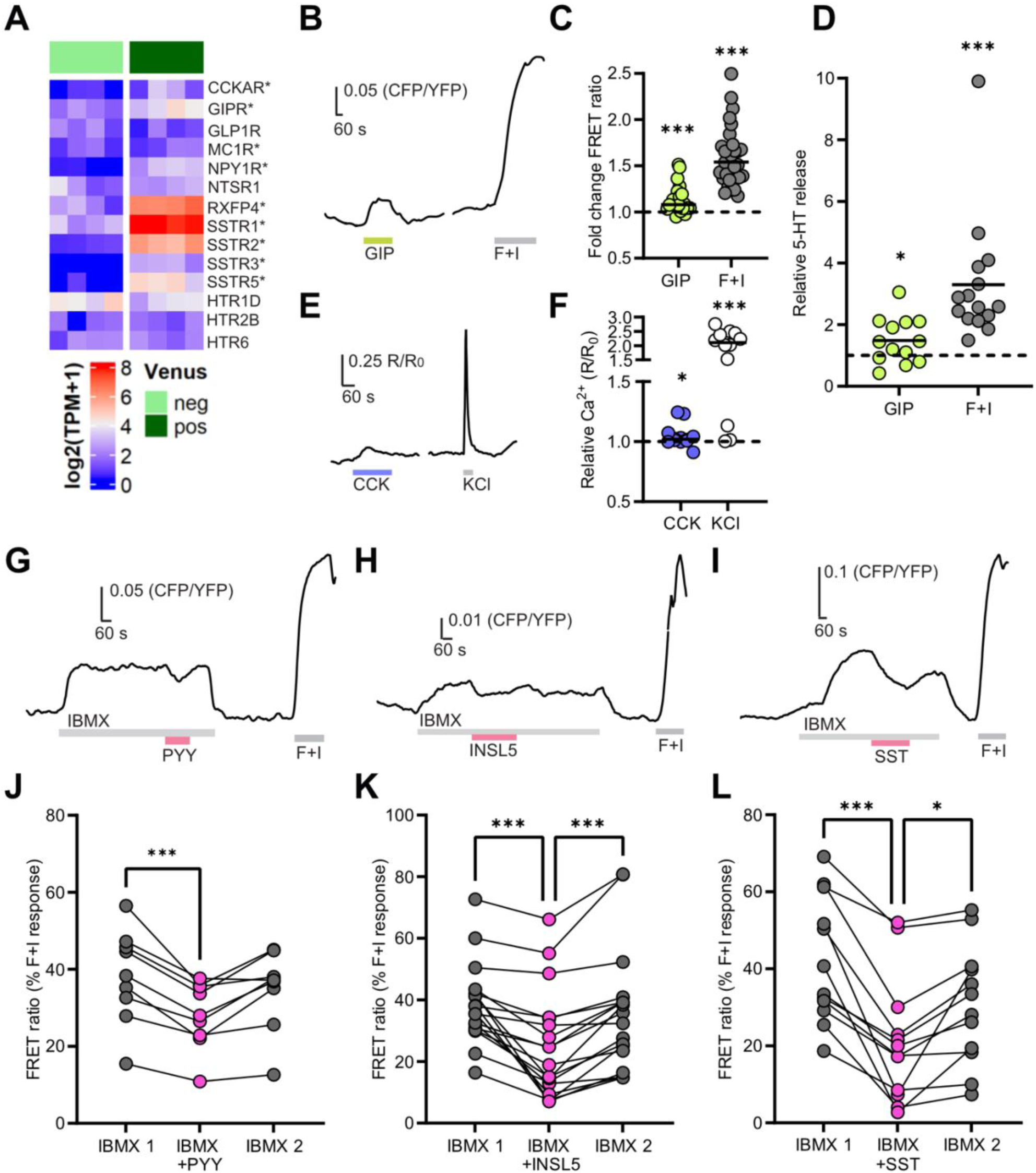
Paracrine activation of human Enterochromaffin cells by gut hormones. A) Heatmap showing expression of selected peptide hormone and monoamine receptors in TPH1 -Venus EC cells (dark green) compared with the Venus-negative population (light green) by bulk RNA sequencing. *depicts significant enrichment in Venus+ population by Wald test (p < 0.05). B) Representative FRET (CFP/YFP) ratio trace of single EC cell being perfused with GIP (100 nM) followed by forskolin + IBMX (F+I, 10 μM and 100 μM, respectively). C) Fold change of FRET ratios recorded as in (B) across n = 22 cells from 3 independent experiments). *** p<0.001 by one sample Wilcoxon test. D) Fold change of supernatant 5-HT concentration in human duodenal TPH1-Venus positive 3D organoids incubated for 1 hour with GIP (100 nM) or Forskolin + IBXM (F+I, 10 μM and 100 μM, respectively). Results are presented relative to concentrations measured in control wells from the same plate (n=13-15 from 3 independent experiments. * p<0.05, *** p<0.001 by one sample t-test. E) Representative Fura2 (340/380 nm) ratio trace of a single EC cell being perfused with CCK (100 nM) and KCl (70 mM). F) Increase in intracellular calcium levels across different cells recorded as in (E), shown as ratio between R (Fura2 ratio during perfusion of stimulus) and R_0_ (Fura2 ratio during perfusion of basal solution). (n =10. 6/10 cells responded to both CCK and KCl, as determined by analysis of z-scores). * p<0.05, *** p<0.001 by one sample Wilcoxon test. G-I) Representative FRET traces of TPH1-Epac-S-H187 EC cells perfused with IBMX (100 μM), in the presence or absence of (G) PYY (100 nM), (H) INSL5 (100nM) and (I) SST-14 (100 nM). Forskolin (10 μM) and IBMX (100 μM) were used as positive control (F+I). J-L) Responses of individual cells recorded as in (G-I). Ratios in IBMX in the presence or absence of PYY (J), INSL5 (K) and SST (L) are expressed as percentage of the forskolin + IBMX maximum response recorded in the same cell (n = 9-18 cells each, from 3-6 independent experiments). * p<0.05, *** p<0.001 by Friedman test with post hoc Dunn’s multiple comparisons test.

### Human Enterochromaffin Cells are Nutrient Sensors

We identified differential expression in human duodenal EC cells of several G-protein coupled receptors (GPCRs) that could respond to nutrients and luminal factors after feeding (Figure 4A), including the olfactory receptors *OR51E1* and *OR51E2*, the short-chain fatty acid (SCFA) receptor *FFAR2*, the 2-acylglycerol receptor *GPR119*, the aromatic amino-acid receptor *GPR142*, and the bile acid receptor *GPBAR1*. The calcium sensor *CASR*, which plays a role in amino acid sensing in GIP-expressing cells^22^, and the long-chain fatty acid receptor *FFAR1* were only detected at very low levels in EC cells, whereas *FFAR4* was detected at higher levels but not differentially expressed.

**Figure 4.**
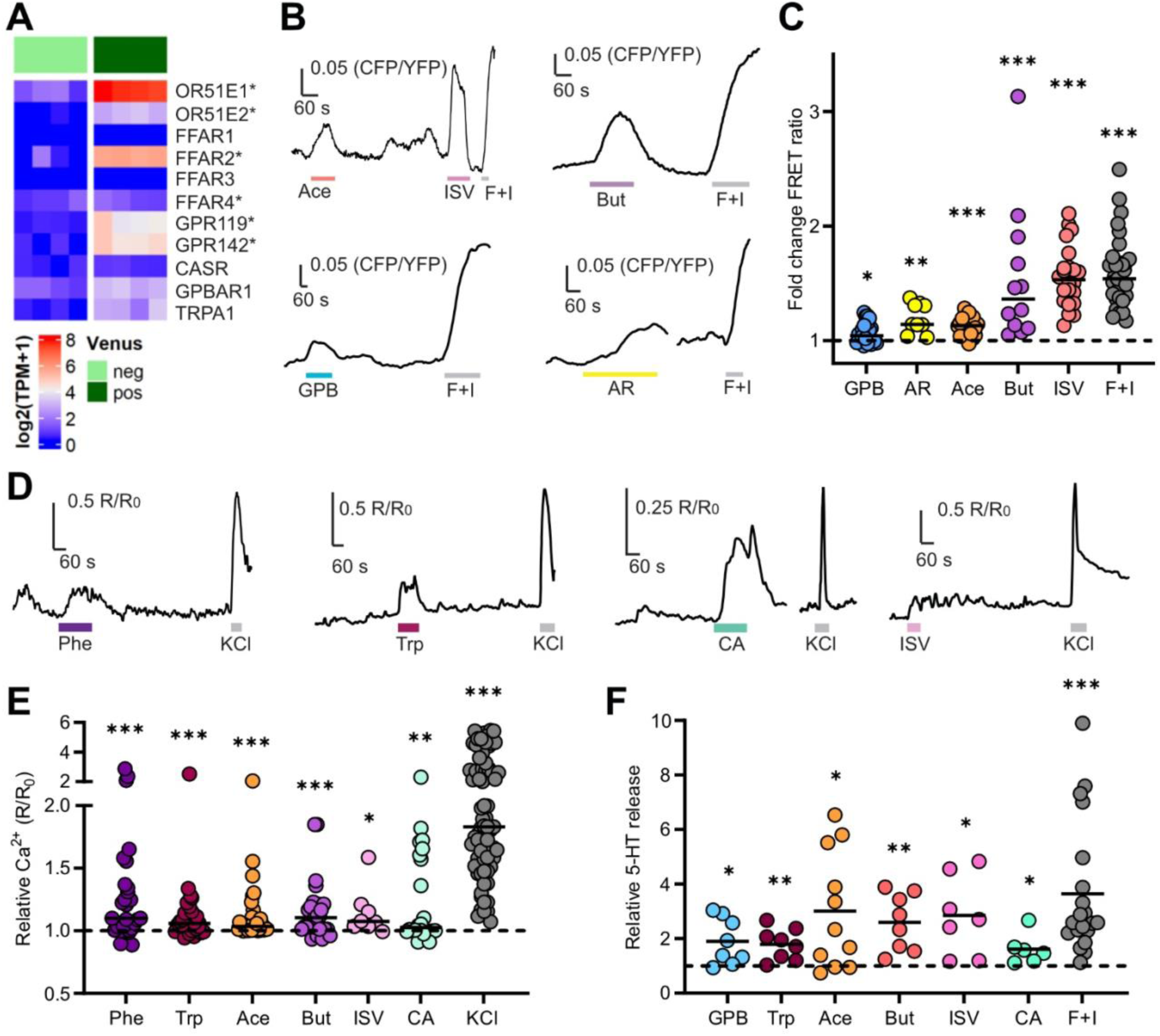
Human Enterochromaffin cells are nutrient sensors. A) Heatmap showing expression of selected nutrient-sensing receptors in TPH1-Venus EC cells (dark green) compared with the Venus-negative population (light green) by bulk RNA sequencing. *Denotes significant enrichment in Venus+ population by Wald test (p < 0.05). B) Representative FRET (CFP/YFP) ratio traces of single EC cells being perfused with acetate (Ace, 200 μM), isovalerate (ISV, 200 μM), butyrate (But, 2 mM), GPBAR-A (GPB, 3 μM), AR231453 (AR, 100 nM) and Forskolin + IBMX (F+I, 10 μM and 100 μM, respectively). C) Fold change of FRET ratio across different cells recorded as in (B). (n = 9-22 cells from 1-3 independent experiments. * p<0.05, ** p<0.01, *** p<0.001 by one sample Wilcoxon test. D) Representative Fura2 (340/380 nm) ratio traces of single EC cells being perfused with phenylalanine (Phe, 20 mM), tryptophan (Trp, 20 mM), cinnamaldehyde (CA, 300 μM), isovalerate (ISV, 200 μM) and KCl (70 mM). E) Data from individual cells recorded as in (D), and shown as ratio between R (Fura2 ratio during perfusion of stimulus) and R_0_ (Fura2 ratio during perfusion of basal solution). (n = 8-62 cells from 3 to 8 independent experiments). * p<0.05, ** p<0.01, *** p<0.001 by one sample Wilcoxon test. F) Fold change of supernatant 5-HT concentration in human duodenal TPH1-Venus positive 3D organoids incubated for 1 hour in the presence of GPBAR-A (GPB, 3 μM), isovalerate (ISV, 200 μM), acetate (Ace, 200 μM), butyrate (But, 1 mM) tryptophan (Trp, 20 mM), cinnamaldehyde (CA, 300 μM) and Forskolin + IBXM (F+I, 10 μM and 100 μM, respectively). Results are presented relative to concentrations measured in control wells from the same plate * p<0.05, ** p<0.01, *** p<0.001 by one sample t-test (n=6-20 from 3 independent experiments).

Functionally, the bile acid receptor agonist GPBAR-A (3 μM), the GPR119 agonist AR231453 (100 nM) and the OR51E1 agonist isovalerate (200 μM)^29^ significantly increased intracellular cAMP in EC cells (FRET changes: 1.04 (0.99-1.15), p=0.01; 1.14 (1.09-1.32), p=0.004; and 1.53 (1.34-1.67), p<0.001, respectively), consistent with the known Gs coupling of these receptors (Figure 4B,C). Acetate is an agonist of both FFAR2 (Gi/q-coupled) and OR51E2 (Gs-coupled)^30^, and our results showed that acetate (200 μM) induced an increase in both cAMP (FRET change 1.13 (1.06-1.20), p<0.001; Figure 4B,C) and Ca^2+^ (1.04 (1.01-1.11) Fura2 R/R_0_, p<0.001; 22/28 cells responded) in human EC cells (Figure 4D,E). Butyrate (1mM), an agonist for FFAR2 and OR51E1, elicited large elevations in Ca^2+^ (1.11 (1.01-1.21) Fura2 R/R_0_, p<0.001; 7/23 cells responded) and cAMP (FRET change: 1.36 (1.12-1.85), p<0.001) (Figure 4B-E). GPBAR-A, acetate, butyrate and isovalerate all induced significant increases in 5-HT release compared to baseline (1.9±0.3, p=0.02; 3.0±0.6, p=0.01; 2.6±0.4, p=0.003; and 2.9±0.5 fold, p=0.015, respectively) (Figure 4F).

We next tested the effects of aromatic amino acids and found that tryptophan (20 mM) and phenylalanine (20 mM) both triggered elevation of intracellular Ca^2+^ in EC cells (Fura2 R/R_0_: tryptophan 1.06 (1.00-1.15), p<0.001, with 17/28 cells responding; phenylalanine 1.10 (1.00-1.36), p<0.001, with 14/29 cells responding) (Figure 4D,E). Tryptophan induced a significant increase in 5-HT release (1.8±0.2 fold, p=0.006) (Figure 4F). These responses are likely mediated by GPR142, as the calcium sensing receptor CASR was expressed only at very low levels in EC-cells, but we cannot exclude other yet to be identified receptors in aromatic amino acid sensing. Finally, the irritant receptor *TRPA1* previously identified in mouse EC cells^2, 3, 31^ was also enriched in human EC cells (Figure 4A). Consistent with the expression data, the TRPA1 agonist cinnamaldehyde (CA, 300 μM) induced an increase in intracellular Ca^2+^ (1.03 (1.00-1.62) Fura2 R/R_0_, p=0.007; 10/23 cells responded) and 5-HT release (1.6±0.2 fold, p=0.04) (Figure 4D-F).

### Nutrient-induced excitability changes in Human Enterochromaffin Cells

Electrical excitability in different types of EEC, including EC cells, has been extensively studied^3, 17, 24, 25, 32^, although there are no previous reports of the electrophysiological characteristics of human EC cells. Human EC cells differentially expressed the voltage-gated Na^+^ channel *SCN3A* (Nav1.3), voltage-gated Ca^2+^ channels *CACNA1H* (T-type / Ca_v_3.2), *CACNA1A* (P/Q-type / Ca_v_2.1), *CACNA1C* (L-type / Ca_v_1.2), as well as the TRP channels *TRPC7* and *TRPM2* (Figure 5A). The mechanosensitive ion channel *PIEZO1* was expressed at similar levels in both EC cells and non-fluorescent control cells, whereas *PIEZO2* was not detected in either (Supplementary Figure 1). We also found enrichment in EC cells of the hyperpolarisation activated non-selective cation channels *HCN4* and *HCN3*, the cyclic nucleotide gated channel *CNGA3*, and *KCNJ3* and *KCNJ6* (Kir3.1/Kir3.2) which together form a G-protein activated inwardly rectifying potassium channel (Figure 5A).

**Figure 5.**
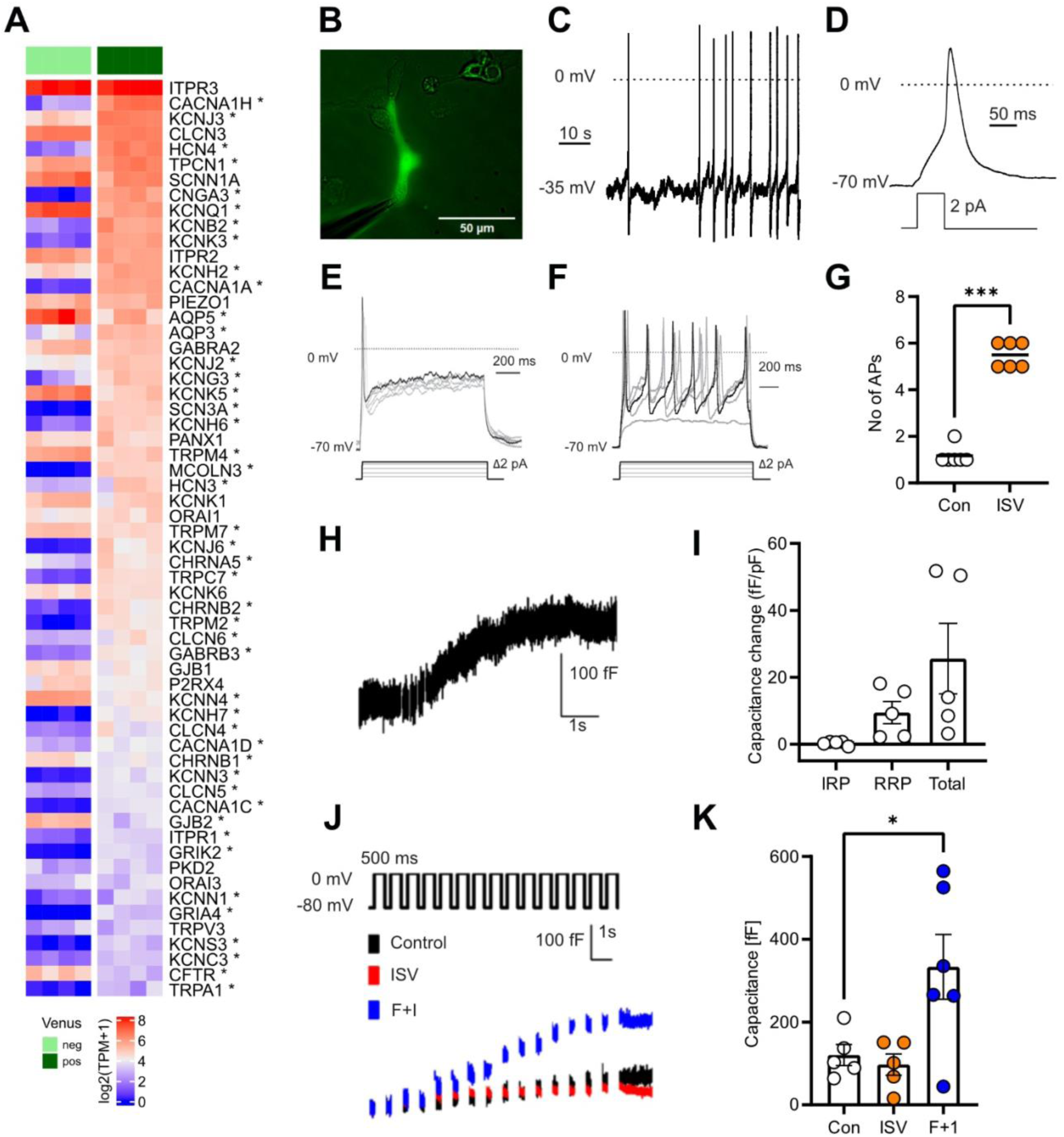
Electrophysiological properties of human Enterochromaffin cells. A) Heatmap showing transcriptomic expression of top 60 ion channels from TPH1+ (dark green) and negative (clear green) populations by bulk RNA sequencing, sorted by decreasing expression in the TPH1+ population. *Represents genes differentially expressed in Venus+ population by Wald test (p < 0.05). B) Example image of human TPH1-Venus+ cell patch clamp experiment. C) Example trace of spontaneous electrical activity of human TPH1-venus+ cell recorded in perforated patch current clamp mode. D) Example trace of action potential from human TPH1-venus+ cell induced by current injection. Prior to the test pulse, the cell was held at −70mV by current injection. E,F) Example traces of current-induced action potentials (1 second pulse) from a single human TPH1-Venus+ cell recorded in control buffer (E) and in the presence of 200 µM isovalerate (F). Cells were held at −70 mV by current injection prior to delivery of the pulse protocol indicated. G) Individual data for cells recorded as in (E,F) showing number of action potentials (AP) triggered by the 30pA injection pulse, in control buffer and in the presence of 200 µM isovalerate (ISV) during the 1 second current injection. ***p<0.001 by paired t-test. H) Example trace of provoked vesicular release (exocytosis) from a human TPH1-Venus+ cell. I) Averaged provoked vesicular release (exocytosis) from human TPH1-Venus+ cells, showing normalized immediate release pool (IRP), readily released pool (RRP) and total release, calculated as indicated in the methods. J) Representative traces of voltage-induced capacitance changes in control solution (Con), 200 µM isovalerate (ISV) and Forskolin + IBMX (F+I; 10 μM and 100 μM, respectively). K) Combined data of voltage-induced capacitance changes from cells recorded as in (J). *p<0.05 by one way ANOVA with post hoc Holm-Šídák’s multiple comparisons test.

We recorded the electrical activity of single human EC cells from 2D organoid cultures (Figure 5B) using perforated patch clamp recordings and found that 9/9 cells fired spontaneous action potentials (resting membrane potential average −67.9±1.9 mV, n=9) (Figure 5C). Properties of evoked action potential were similar to those previously reported in human K-cells^22^ and L-cells^33^ (action potential threshold −23.5±3.2 mV, overshoot 35.5±3.1 mV and half-width 20.17±1.0 ms, n=9; Figure 5D). Acute treatment with isovalerate induced an increase in action potential firing elicited by 1-sec long depolarising current injections (1.7±0.4 vs 5.5±0.5 events per 1-sec pulse, n=6. Paired t-test, p<0.05) (Figure 5E-G).

Whole cell voltage-clamp electrophysiological capacitance recordings were used to assess depolarisation-induced exocytosis by human EC cells. To dissect distinct functionally defined secretory vesicle pools in human EC cells, we employed a depolarization protocol well-characterized for the study of exocytosis in adrenal chromaffin cells^34^. Human EC-cells had an average initial capacitance (a measure of cell size) of 10.8±0.9 pF (n=5). An example capacitance trace is shown in Figure 5H, and mean data from the 5 capacitance traces recorded using this protocol are shown in Figure 5I. We measured the “immediately release pool” (IRP) in response to a series of 6 short 10ms depolarizations (3.5 ± 2.6 fF per cell (0.28 ± 0.27 fF/pF)), the “readily releasable pool” (RRP) after four additional long 100ms depolarisations (91 ± 27 fF per cell (9.5 ± 3.3 fF/pF)), and the mean total vesicular release, determined as the change in membrane capacitance from zero to seven seconds (240 ± 89 fF per cell (25.6 ± 10.6 fF/pF)) (Figure 5I). Our findings indicate that human EC cells have a large functional vesicle pool to support exocytosis similar to other endocrine cells^34^.

Next, we investigated if chemical stimuli could enhance exocytosis triggered by membrane depolarisation. For this, we measured EC cell capacitance in response to a train of 500ms depolarising pulses from −80mV to 0mV in either regular extracellular solution or in the presence of 200 μM isovalerate or forskolin+IBMX (10 μM and 100 μM, respectively) (Figure 5J,K). Whereas forskolin+IBMX induced an increase in pulse-evoked exocytosis compared with control (48.7±16.5 fF vs 26.8±9.9 fF, respectively. ANOVA, *p<0.05, n=5-6), isovalerate did not (19.1±4.8 fF, n=5).

## DISCUSSION

Human duodenal organoids expressing fluorescent proteins and sensors in EC cells provide a method to characterize the mechanisms underlying gut 5-HT release – a question that has previously been hampered by a shortage of representative in vitro models and difficulties in measuring acute 5-HT release in plasma due to the high background storage of 5-HT in platelets. Like mouse EC cells and other human EECs, human EC cells were electrically active and exhibited enhanced exocytosis, measured by capacitance recordings, upon membrane depolarization and elevation of cAMP. Transcriptomic analysis of purified organoid-derived human duodenal EC cells identified expression of a range of GPCRs, which were functionally validated by fluorescence-based recordings of cytoplasmic Ca^2+^ and cAMP levels, and by ELISA-based measurements of 5-HT secretion.

Although *TPH1* was the most highly enriched hormonal marker in human EC cells, the cells also exhibited enriched mRNA expression and detectable peptide levels of other gut hormones including SST, CCK and PYY, although not of motilin (MLN) or glucose-dependent insulinotropic polypeptide (GIP). This is consistent with previous reports that EC cells show transcriptomic overlap with other EEC populations^35, 36^. However, our recent single cell RNA sequencing analysis of chromogranin-A positive cells purified from human duodenal CHGA-Venus organoids, returned distinct EEC clusters showing only limited overlap between EC cells and cell populations expressing *SST*, *CCK* and *PYY*, despite identifying co-expression of different hormones in a substantial proportion of cells^37^. Consistent with studies on murine EC cells we found enriched expression of tachykinin genes in human EC cells, however, unlike murine cells which predominantly express *Tac1*, we found both *TAC3* and *TAC1* in human cells, with *TAC3* expression being higher than that of *TAC1*. *TAC1* encodes Neurokinin A (NKA) and Substance P, whilst *TAC3* encodes Neurokinin B (NKB). Interestingly, a recent study reported that polymorphisms in *NK2R* (the cognate receptor for NKA > NKB) are associated with body weight and glucose control in humans, and that agonism of NK2R in mouse models resulted in reduced food intake, lower body weight and loose stools^38^. The physiological role of EC-derived tachykinin peptides in humans is not clear, although substance P is implicated in pain perception in the intestine^39^.

Amongst the GPCRs identified and characterized in human EC cells were members of the adrenergic receptor family, responsive to adrenaline and noradrenaline. Adrenergic stimulation of 5-HT release from human organoid-derived EC cells was predominantly mediated by β-adrenergic receptors, as also previously reported from measurements of portal blood 5-HT in cats after stimulation of sympathetic fibers carried in the vagus nerve^40^. Human EC cells exhibited small Gi-dependent calcium responses to α2 adrenoceptor agonism, mirroring findings in mouse EC cells that have been attributed to downstream Trpc4 channel activation^3^, but in human EC cells Ca^2+^ responses to clonidine were accompanied by a reduction in cAMP levels and no net stimulation of 5-HT secretion. β-Adrenergic agonism with isoproterenol, by contrast, increased human EC cell Ca^2+^ and cAMP levels and 5-HT secretion. This appears to differ from murine EC cells which have been reported not to express detectable β-adrenergic receptor mRNA transcripts or to exhibit responses to β-adrenergic agonism^3^, although our recent scRNAseq analysis of murine EEC cells from different intestinal regions did detect β-adrenergic receptor expression in a number of EEC-clusters, including EC cells^41^.

Human EC cells exhibited detectable expression receptors for several gut hormones, including the Gs coupled receptors *GIPR* and *GLP1R*, the Gq coupled receptors *CCKR1* and *NTSR1*, and the Gi coupled receptors *NPY1R*, *RXFP4*, *SSTR1,2,3* and 5, suggesting paracrine modulation of EC cells by other neighbouring EECs. A number of these GPCRs were validated functionally; agonism of GIPR increased cAMP and enhanced 5-HT release, whilst PYY, SST and INSL5 lowered cAMP in EC cells from human organoid cultures. Previous studies in proximal and distal mouse intestine also found evidence of paracrine communication between L-cells and EC cells^11, 42^, but as PYY, INSL5 and GLP-1 are co-released by L-cells in response to nutrients^43–46^, it is surprising that these hormones have opposing effects on EC cells, with GLP-1 being stimulatory and PYY and INSL5 being inhibitory. Stimulation of mouse EC cells by GLP-1 was reported previously^11^, and *in vitro* studies using a human epithelial cell line also reported that INSL5 decreased cAMP levels and inhibited 5-HT release^21^. The *in vivo* balance of stimulatory and inhibitory interactions between L-cells and EC cells thus remains unclear, and direct effects of INSL5 on EC cells do not appear to account for observations that INSL5 administration or stimulation of INSL5-expressing L-cells induces a 5-HT-dependent increase in colonic motility in mice, as this would require stimulation not inhibition of EC cells by INSL5^42, 47^. It is possible that pro-motility effects of INSL5 *in viv*o require 5-HT-dependent signalling in the enteric nervous system, rather than release from EC cells.

Although previous studies suggested that EC cells do not express high levels of nutrient-sensing GPCRs, we found expression and functional activity of GPBAR1, GPR119 and GPR142 in human EC cells, potentially enabling the cells to respond to stimuli such as bile acids, monoacylglycerols and aromatic amino acids^22^. Another recent study of human organoid-derived EECs also investigated associations between GPCR expression and 5-HT (and other hormones) release using a receptor knockout approach^48^. The authors reported effects of receptor agonists or antagonists on 5-HT secretion during 16 h incubations and compared 5-HT in the supernatant in WT and knockout organoids to a control stimulation with forskolin and IBMX. Whilst several GPCR agonists stimulated a degree of 5-HT release, knockouts of *FFAR2, CASR, GPR142, GPR119* or *GLP1R* did not significantly affect responses to their respective agonists. Exceptions were FFAR4, GPR19 and MC1R, which we have not further explored in our study, but our functional data presented here would support roles for FFAR2, GPR142, GPR119 and GIPR, whilst CASR seems not to be expressed to sufficient levels in EC cells. It is important to highlight that long-term exposure to these agonists in secretion experiments might additionally alter gene expression or activate paracrine signalling from neighbouring EECs that will ultimately affect 5-HT secretion. Our functional assays additionally confirm that receptor agonists can directly activate single human EC cells via intracellular Ca^2+^ and/or cAMP.

When we compared human EC cells with other EEC populations using data from CHGA-Venus organoids^37^, we found that expression levels of *GPBAR1*, *GPR119* and *GPR142* in EC cells were similar to those in duodenal I-cells (CCK) and K-cells (GIP), whereas expression of *CASR* and *FFAR1* were notably lower in EC cells than in other EECs from the same intestinal region. Low expression of *FFAR1* is unusual for EECs^22, 24^ and suggests a specific lack of FFAR1-dependent sensing of long chain fatty acids by EC cells. Orally ingested lipids could still, however, potentially stimulate 5-HT release via monoacylglycerols acting on GPR119, and by GPBAR1-dependent action of bile acids present in lipid micelles.

EC cells exhibited high expression of *OR51E1* and *FFAR2*, which are responsive to short chain and branched chain fatty acids produced by microbial fermentation. The olfactory receptors OR51E1 and OR51E2, and their mouse homologues Olfr558 and Olfr78, have been identified previously in mouse and human EECs, particularly in the lower GI tract where microbiota levels are high. The observed Ca^2+^ responses to acetate and butyrate in human EC cells are likely attributable to the Gi/q coupled receptor FFAR2, and the cAMP responses to butyrate and isovalerate likely reflect the Gs coupling of OR51E1. As OR51E1 is not considered sensitive to acetate, the observed acetate-triggered cAMP response may reflect OR51E2 activity. We were surprised, and cannot explain, that although isovalerate increased cAMP levels and 5-HT secretion, it did not enhance EC cell secretion measured by capacitance recordings; one possible explanation is that cAMP responses to forskolin + IBMX are larger and more sustained than those to isovalerate in the vicinity of secretory vesicles. Alternatively prolonged exposure to isovalerate might result in desensitisation of the signals enhancing capacitance responses to strong depolarising pulses, consistent with a detectable decline in the cAMP response during the short exposure shown in Figure 4B; however, the increased action potential frequency in response to depolarising pulses was observed after exposure to isovalerate for several minutes, which might contribute to the increase in serotonin secretion during a 1 h exposure.

### Conclusions

Human EC cells are modulated by a range of intestinal chemical stimuli, including sympathetic tone, paracrine actions of other gut hormones, ingested nutrients and microbial metabolites – in addition to their previously-demonstrated responsiveness to mechanical forces. As EC cell activity contributes to the regulation of intestinal motility, conveys signals related to intestinal pain, and initiates pathways that regulate appetite, understanding the mechanisms controlling human EC cells provides opportunities to intervene therapeutically in a range of clinical disorders of the gut brain axis.

**Table 1.** Drugs used for secretions, perfusions, imaging and electrophysiological studies.

| Reagent | Description | Source | Catalogue number |
| --- | --- | --- | --- |
| Adrenaline | Adrenoceptor agonist | Sigma-Aldrich | E4250 |
| AR231453 | GPR119 agonist | Sigma-Aldrich | SML1883 |
| Clonidine hydrochloride | α2-adrenergic agonist | Tocris | 0690/100 |
| D-(+)-Glucose | Nutrient | Merck Life Science UK Limited | G7528-250G |
| Exendin-4 | GLP-1R agonist | Cambridge Bioscience | 4027457.1000 |
| Fluoxetine hydrochloride | SERT inhibitor | Fluorochem | F375072 |
| Forskolin | Adenylate cyclase activator | Sigma-Aldrich | F6886 |
| Fura2-AM | Cell permeable Ca<sup>2+</sup> indicator | Invitrogen | F1221 |
| GPBAR-A | G-protein bile acid receptor 1 agonist | Sigma-Aldrich | SML1207 |
| Human GIP | GIPR agonist | Cambridge Bioscience | AS-61226-05 |
| Human PYY 3-36 | NPY Y2 receptor agonist | Tocris | 6288 |
| Human CCK | CCRAR agonist | Phoenix Pharmaceuticals | 069-27 |
| IBMX (3-isobutyl-1-methylxanthine) | Phosphodiesterase inhibitor | Sigma-Aldrich | I7018 |
| INSL5, short A- & B-Chains | Rxfp4 agonist | Phoenix Pharmaceuticals | 035-70A |
| Isovaleric acid | OR51E1 agonist | Acros Organics | 156690025 |
| L-Phenylalanine | Aromatic amino acid | Sigma | P5482-25G |
| L-Tryptophan | Aromatic amino acid | Sigma-Aldrich | T0254 |
| Noradrenaline | Adrenoceptor agonist | Sigma-Aldrich | N5785 |
| Sodium acetate trihydrate | Short-chain fatty acid | Sigma-Aldrich | 6131-90-4 |
| Sodium butyrate | Short-chain fatty acid | Sigma-Aldrich | B5887 |
| Somatostatin 14 | SSTR1-5 agonist | Generon Ltd | AS-24277 |

## ACKNOWLEDGEMENTS

We thank the MRL Genomics and Transcriptomics Core, the Core Biochemical Assay Laboratory (CBAL), the Flow Cytometry Core at CIMR, CRUK Cambridge Institute Genomics Core and Addenbrooke’s Tissue Bank and Esmira Mamedova for help with imaging and electrophysiology training. For the purpose of open access, the author has applied a Creative Commons Attributions (CC BY) license to any Author Accepted Manuscript version arising from this submission.

## Abbreviations

cAMP: cyclic adenosine monophosphate
CCK: cholecystokinin
5-HT: 5-hydroxytryptamine
EC: Enterochromaffin
EEC: enteroendocrine cell
FRET: Fluorescence resonance energy transfer
GIP: glucose-dependent insulinotropic peptide
GPCR: G-protein coupled receptor
IBMX: 3-isobutyl-1-methylxanthine
INSL5: insulin-like petide 5
PYY: peptide YY
SCFA: short chain fatty acid
SST: somatostatin.

## Transcriptomic profiling

RNA sequencing data will be deposited in the National Center for Biotechnology Information-Gene Expression Omnibus (NCBI GEO) repository (accession number to follow).

## Peptidomics

Data will be deposited to the ProteomeXchange Consortium via the PRIDE partner repository (accession number to follow).

## Author contributions

Study conception and design: CA, NG, EM, CI, FR, FG. Data acquisition, analysis and/or interpretation: CA, NG, EM, JRQ, AD, CS, EO, MSH, MT, TL, MH, RBB, RK, AS, CI, FR, FG. Manuscript drafting: CA, FR, FG. Final manuscript approval, accountability for all aspects of the work: CA, NG, EM, JRQ, AD, CS, EO, MSH, MT, TL, MH, RBB, RK, AS, CI, FR, FG.

## Data transparency

Study materials and raw data will be made available upon application to the senior authors.

