## Supplementary Figure 1 for "Mechanisms of Activation and Serotonin release from Human Enterochromaffin Cells"

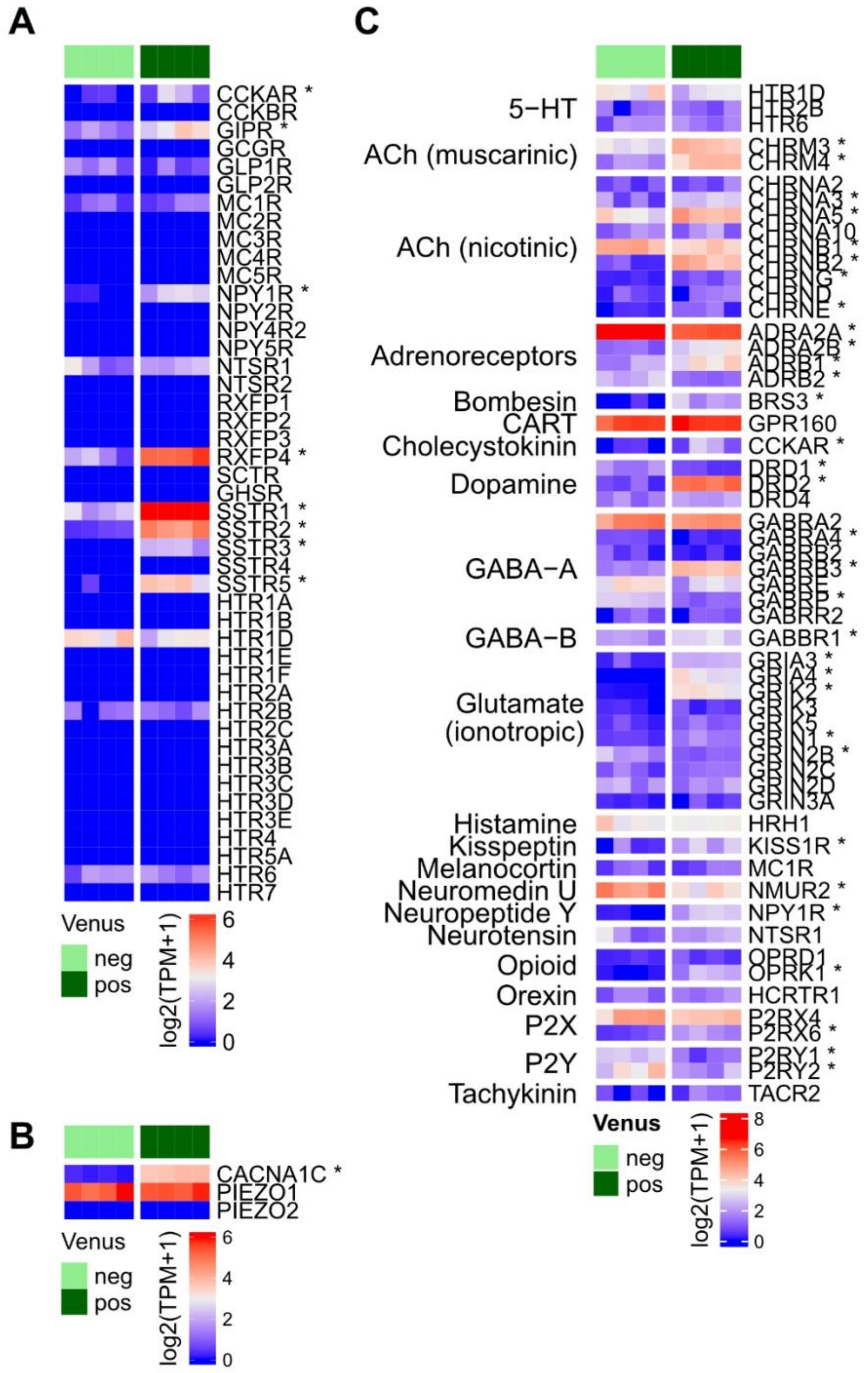


Transcriptional expression of hormone receptors, ion channels and neurotransmitter receptors from TPH1+ (dark green) and negative (clear green) populations by bulk RNA sequencing. A) Heatmap showing expression of hormone/peptide receptors in TPH1-Venus

EC cells. B) Heatmap showing expression of Piezo1 and Piezo2 mechanosensitive ion channels in comparison to the differentially enriched voltage-gated calcium channel CACNA1C in TPH1-Venus EC cells. C) Heatmap showing the expression of neurotransmitter receptors in TPH1-Venus EC cells. *Genes significantly enriched in Venus+ population by Wald test (*$p < 0.05$).
